# An evaluation of multi-excitation-wavelength standing-wave fluorescence microscopy (TartanSW) to improve sampling density in studies of the cell membrane and cytoskeleton

**DOI:** 10.1101/2020.11.26.396622

**Authors:** Jana K. Schniete, Peter W. Tinning, Ross C. Scrimgeour, Gillian Robb, Lisa S. Kölln, Katrina Wesencraft, Nikki R. Paul, Trevor J. Bushell, Gail McConnell

## Abstract

Conventional standing-wave (SW) fluorescence microscopy uses a single wavelength to excite fluorescence from the specimen, which is normally placed in contact with a first surface reflector. The resulting excitation SW creates a pattern of illumination with anti-nodal maxima at multiple evenly-spaced planes perpendicular to the optical axis of the microscope. These maxima are approximately 90 nm thick and spaced 180 nm apart. Where the planes intersect fluorescent structures, emission occurs, but between the planes are non-illuminated regions which are not sampled for fluorescence. We evaluate a multi-excitation-wavelength SW fluorescence microscopy (which we call TartanSW) as a method for increasing the density of sampling by using SWs with different axial periodicities, to resolve more of the overall cell structure. The TartanSW method increased the sampling density from 50% to 98% over seven anti-nodal planes, with no notable change in axial or lateral resolution compared to single-excitation-wavelength SW microscopy. We demonstrate the method with images of the membrane and cytoskeleton of living and fixed cells.

## Introduction

Confocal laser scanning microscopy (CLSM) [1] and multi-photon laser scanning microscopy [2] are now widely used for 3D cell imaging. Both provide optical sectioning by elimination of the contribution of out-of-focus parts of a 3D specimen, but both have limited axial resolution. Improved resolution is needed to study many processes in cell biology, such as the nanoscale organisation of the membrane [3] or the nanomechanics of the actin cytoskeleton in cells [4].

To improve the spatial resolution of a cell image, CLSM has also been combined with other imaging methods, such as atomic force microscopy (AFM) [5,6]. This multi-modal approach simultaneously provides mechanical data from the cell, but instruments are complex and the acquisition time for both 3D AFM and CLSM data, is very long compared to many dynamic processes in the cell including reorganization of the actin cytoskeleton [7].

True improvements in resolution surpassing the Abbe-Rayleigh limit have now been achieved by a number of methods. Methods such as photoactivated localization microscopy (PALM) and Stochastic Optical Reconstruction Microscopy (STORM) achieve a lateral resolution of around 10-50 nm by computational localization of the source fluorophore within the point-spread function (PSF) of the objective lens [8]. In stimulated emission depletion (STED) microscopy the fluorescence emission is limited to a tiny central volume of the PSF through generation of a doughnut-shaped STED beam [9–11]. These methods have allowed tracking of individual actin molecules, and the study of actin polymerisation areas in dendritic cells [12], as well as imaging the two layers of actin at the leading edge of the cell [13]. These super-resolution techniques are, however, hampered by limited probe availability [14]. In addition, the long image acquisition times for both 2D and 3D super-resolution imaging can be prohibitive, particularly for live cell studies, and these methods are usually suitable for imaging only a very small field of view due to the requirement of high magnification, high numerical aperture objective lenses [15,16]. Total internal reflection fluorescence (TIRF) microscopy can improve the axial resolution to 70 nm, thus overcoming the diffraction limit, but this technique is limited to 2D imaging of the basal cell membrane[17] as is the comparable method supercritical illumination microscopy [18].

An improvement in both lateral and axial resolution of up to a factor of 2 can be achieved by the use of Structured Illumination Microscopy (SIM) [19,20], which is an intrinsically slow and computationally intensive method, but has been made faster by optomechanical design improvements.

Conventional standing-wave (SW) fluorescence microscopy uses a single wavelength to excite fluorescence from the specimen and is an affordable and effective method to increase axial resolution. It requires only that a reflector is placed close to the specimen when using an epifluorescence microscope, although more complex means may also be used to create the necessary counterpropagating coherent beams of the same wavelength [21,22]. This method improves the axial resolution but not the lateral resolution. The SW is comprised of alternating nodal and anti-nodal planes that are oriented orthogonally to the optical axis of the microscope. The anti-nodal maxima, in which fluorescence can be excited in the specimen, are separated along the optical axis by λ/2n and are λ/4n thick, where λ is the wavelength of light and n is the refractive index of the cell [22,23]. The λ/4n thick illumination plane determines the axial resolution of the image, and is typically 90 nm thick (at the full-width at half-maximum intensity limit).

One problem is that even with objectives of the highest available numerical aperture, the PSF extends to 600 nm axially, so one image represents the fluorescence signal detected from several anti-nodal maxima, if there is fluorescent material present in the multiple planes. Consequently, the method is best suited to study sparse cell specimens including stress fibres dissociated from mouse embryo fibroblast 3T3 cells [21], and fluorescently-labelled cell membranes, where the regularly-spaced anti-nodal maxima intersect with the membrane leading to an emission pattern that resembles a contour map. We have recently used SW microscopy to study 3D deformations in the red blood cell membrane at frame rates of up to 100 Hz, with an axial resolution of around 90 nm, using a conventional widefield epifluorescence microscope [24]. SW microscopy has also been demonstrated in confocal laser scanning mode, even though wavefronts that are reflected from the mirror are not strictly parallel to the specimen plane [25].

A limitation in SW microscopy is that unlike in full-field epifluorescence, fluorophores are only detected in the anti-nodal planes. Between planes there is therefore a *terra incognita* where no fluorescence can be detected, which amounts to more than half of the volume of a cellular specimen. Another limitation of SW microscopy is that the image resembles a contour map, but without the marked heights. This introduces ambiguity into the image dataset: for example, it is not possible to confirm that an object in a SW image is convex or concave.

Here, we report a new method called TartanSW that uses a confocal microscope and multiple excitation wavelengths to create multiple SWs for excitation of fluorescence in the cell specimen. In TartanSW, we use more than one excitation wavelength to create multiple SWs of different periodicity in the specimen. Since the anti-nodal planes generated with different wavelengths rarely coincide, the density of sampling is thereby increased and the data gap between the planes that ordinarily appear in single-wavelength-excitation SW microscopy is reduced. This is shown schematically in Figure 1, with an adherent cell on top of a mirror that is illuminated from above, as per an upright microscope. The conventional single-wavelength-excitation method is shown in Figure 1(A) with green anti-nodal planes of thickness λ/4n separated by λ/2n (not shown to scale). The effect of the TartanSW multiwavelength-excitation is shown in Figure 1(B), where two further SW patterns of higher (blue) and lower (red) spatial frequencies are added. Since the anti-nodal planes of different SWs occur with different frequencies and do not regularly occur at the same axial height relative to the mirror, we set out to test here whether the sampling density of the imaged cell could be increased by the introduction of additional SWs at new wavelengths.

**Figure 1:**
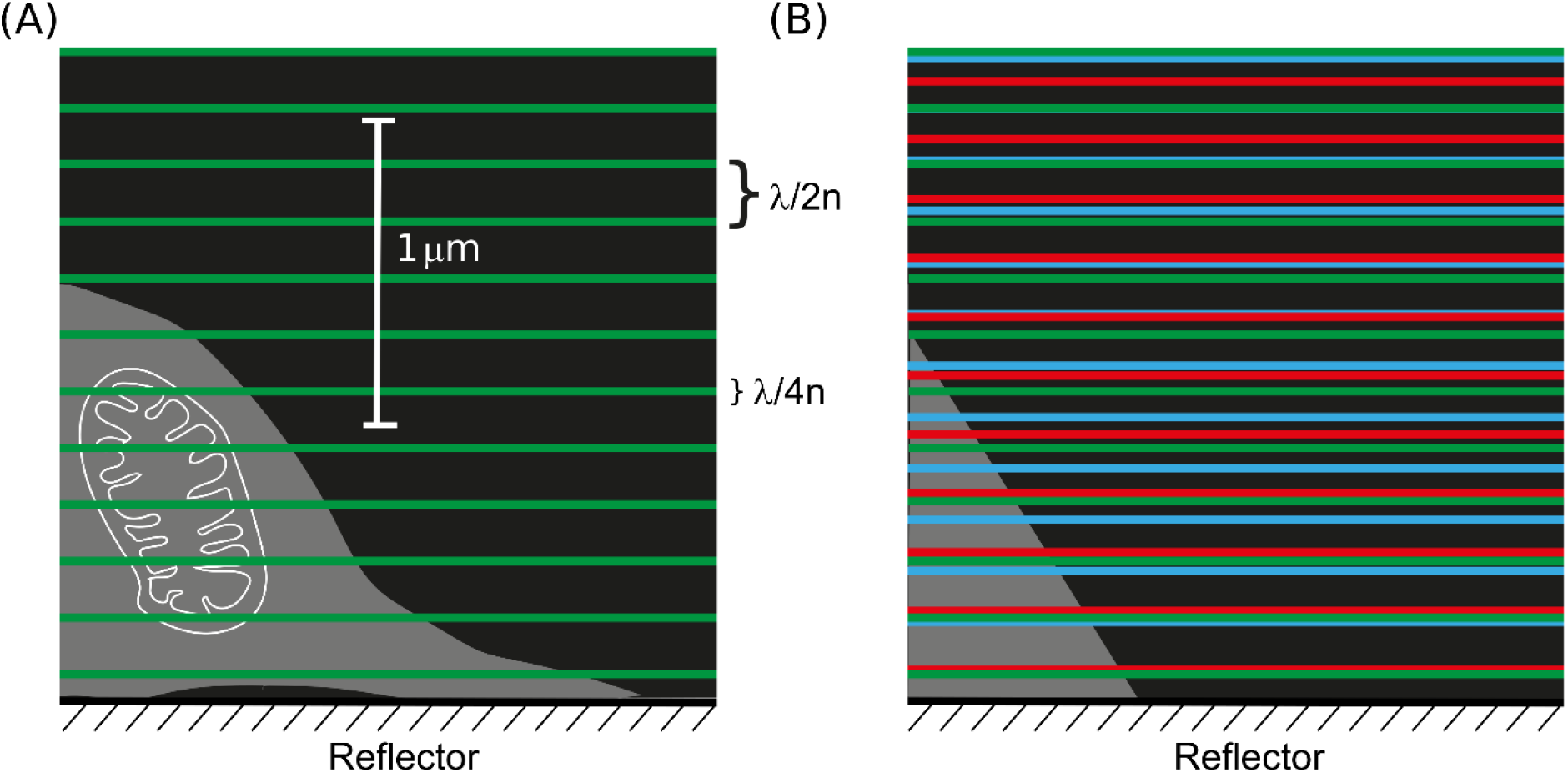
Schematic overview of TartanSW imaging. (A) single-wavelength-excitation SW principle. The cell (mitochondrion in white included for scale) adheres to the mirror, and is illuminated from above, as in an upright microscope. The green lines represent the anti-nodal planes formed by the SW pattern. Each anti-nodal plane has a thickness equivalent to λ/4n, and the spacing between the anti-nodal planes is equivalent to λ/2n, where λ is the respective wavelength and n is the refractive index. (B) multi-wavelength-excitation SW imaging (TartanSW) with blue, green and red lines representing the axial position of the anti-nodal planes from multiple SWs of different frequencies. In both (A) and (B) the anti-nodal plane thicknesses are reduced for clarity but the scaling of spatial frequencies is correct for 488 nm, 514 nm and 543 nm, assigned to false colours blue, green and red respectively. The medium was PBS (n=1.34). For all frequencies, a nodal plane (not marked) lies in the surface of the reflector.

We also predicted that it should be possible to use the relative axial position of the anti-nodal planes occurring at different wavelengths to infer the shape of the cell structure. By using multiple excitation wavelengths, we hypothesised that it would be possible to recognize the order of the SW anti-nodes by their colours and so tell the difference between objects with a positive or negative curvature relative to the anti-nodal planes.

Our method takes advantages of two commonly overlooked but not unrelated properties of current optical microscopy methods and equipment for cell imaging. Firstly, the growing range of fluorescent dyes and photoproteins with broad excitation spectra allows the use of more than one excitation wavelength without the need to create new and sophisticated fluorophores. Secondly, confocal microscopes are routinely fitted with sources of multiple laser lines, giving multiple excitation wavelengths as standard. This combination means that existing instrumentation can be used with our new method, with no adaptation to the microscope hardware or specimen preparation protocol. We call this method TartanSW because of the striking similarity of the coloured fringe patterns seen in our image data to those found in tartan patterned fabrics.

To evaluate whether our proposed TartanSW method would indeed increase the sampling density and whether colour-ordering could be used to infer cell shape, we first performed imaging of a non-biological test specimen, namely a spherical lens surface coated with a thin layer of fluorescent dye [24]. We used a custom MATLAB script to measure the difference in sampling density that results from the TartanSW method compared to single-excitation-wavelength SW microscopy. We then evaluated the TartanSW method using live and fixed cells, labelling the plasma membrane and actin network of mammalian cells with fluorescent probes or common photoproteins, and we explored whether colour-ordering in TartanSW could reduce or remove the ambiguity of cell shape that occurs in conventional SW microscopy.

## Results

### TartanSW imaging of the non-biological test specimen

With the thin spherical fluorescent surface, it proved possible to perform sequential SW imaging to test the basic proposition that sampling density could be increased with the TartanSW method.

Figure 2 shows the single-wavelength-excitation SW images of the lens specimen obtained with (A) 488nm excitation, pseudo-coloured blue, (B) 514 nm excitation, pseudo-coloured green, and (C) 543 nm excitation, pseudo-coloured red. A pseudo-colour RGB merge of images (A)-(C) to create the TartanSW image is shown in 2(D).

**Figure 2:**
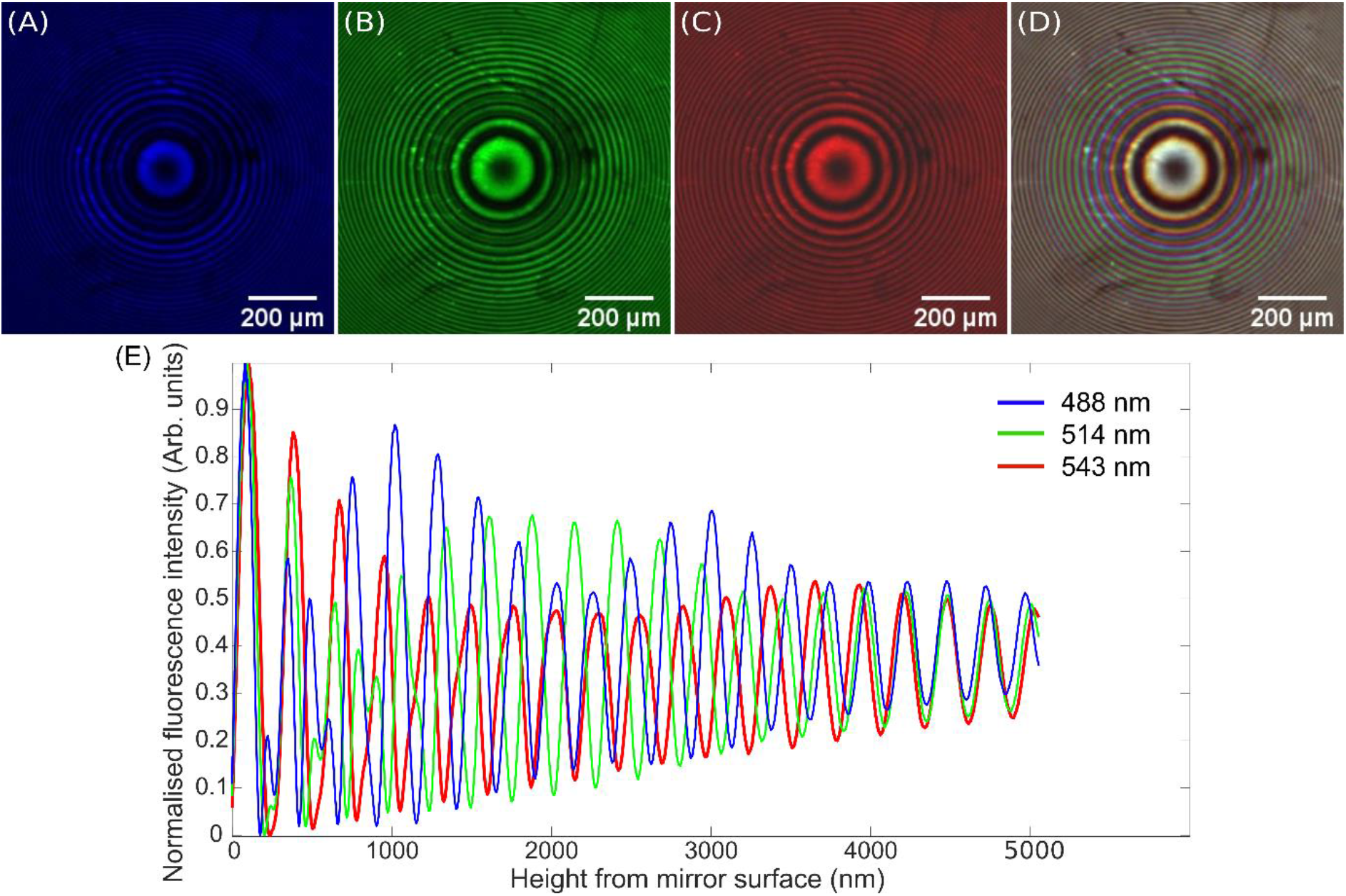
Single-excitation-wavelength SW images of a DiI coated f = 48 mm lens specimen obtained using excitation wavelengths of (A) 488 nm, (B) 514 nm and (C) 543 nm, which are pseudo-coloured blue, green and red respectively, and (D) shows the RGB composite of 2(A)-(C) which is the TartanSW image. The lens specimen was placed on a mirror (Laser 2000) and was imaged using a Leica SP5 confocal microscope with a 5x/ 0.15 N.A. HCX PL FLUOTAR DRY objective. Images were 2048 × 2048 pixels in size, and were averaged over 3 frames at a line speed of 100 Hz, with an emission detection bandwidth of 550-650 nm. The scale bar for images (A)-(D) is 200 μm. (E) Normalized intensity profiles of the lens specimen from (A)-(C) computed using the custom MATLAB code for excitation wavelengths of 488nm (blue), 514 nm (green) and 543 nm (red). As expected, the frequency of the anti-nodal planes increases as the excitation wavelength decreases, and thus the position of the anti-nodal planes for the different wavelengths does not always coincide. By analysis of the full-width at half-maxima of the single-excitation-wavelength intensity profile for 543 nm only, the lens is imaged with a sampling density of 50%. Introducing two further SWs at excitation wavelengths of 488 nm and 514 nm increases this sampling density to 98%.

In Figure 2(D), near-coincident anti-nodal planes near the mirror surface result in a single, white-coloured ring, with colours barely visible at the interior and exterior of this ring, indicative of the small variation in anti-nodal plane separation. This is to be expected because of the nodal plane occurring at the mirror surface. However, as predicted, each of the individual excitation SWs are increasingly out-of-phase with respect to each other as the distance from the mirror increases, forming collectively a dense moiré-like pattern.

These data were analysed using the custom MATLAB script as described above, and the results are presented in 2(E). The sampling density of the single-excitation-wavelength image obtained at 543nm was measured to be 50%. After introducing two additional SWs at excitation wavelengths of 488 nm and 514 nm to create the TartanSW image, the sampling density increased to 98%. A 3D reconstruction of these data, made with MATLAB, is shown in Supplementary Figure 3. Table 1 shows the theoretical and measured anti-nodal plane thicknesses and separations, with the measured values taken from the data shown in Figure 2(E). The theoretical values were calculated using λ/4nm and λ/2n respectively, with λ being the excitation wavelength and a refractive index of n=1.34. The measured values are the FWHM. All measured anti-nodal spacings were consistent with the theoretical values. The measured anti-nodal plane thickness values were in agreement with the theoretical values.

**Table 1.** Theoretical and measured anti-nodal plane thicknesses and separations for the excitation wavelengths used to image the lens specimen coated with DiI. The measured values are taken from the data shown in Figure 2(E). The theoretical values of anti-nodal spacing and anti-nodal plane thickness (FWHM) were taken using λ/2n and λ/4n respectively, with λ being the excitation wavelength and a refractive index of n=1.34.

| Excitation wavelength (nm) | Theoretical anti-nodal spacing (nm) | Measured anti-nodal spacing (nm) | Theoretical anti-nodal plane thickness (nm) | Measured anti-nodal plane thickness(nm) |
| --- | --- | --- | --- | --- |
| 488 | 244 | $239 \pm 7$ | 122 | $122 \pm 4$ |
| 514 | 257 | $254 \pm 3$ | 129 | $136 \pm 1$ |
| 543 | 271 | $270 \pm 6$ | 136 | $139 \pm 1$ |

### TartanSW cell imaging

Figure 3 shows pseudo-coloured cell images obtained using the TartanSW method, each resulting from three different excitation wavelengths applied sequentially. Figure 3(A) shows live MCF-7 cells (female breast cancer cells) stained with the lipophilic stain DiI for the cell membrane, with a cropped and zoomed region of interest (ROI) of 3(A) shown in 3(B). Figure 3(C) shows live SKOV-3 cells (female ovarian cancer cell line) labelled with a different membrane dye, Lipilight 560, with a cropped and zoomed ROI in 3(D). Figure 3(E) shows fixed 3T3 cells (mouse embryo fibroblast cell line) prepared with rhodamine-conjugated phalloidin to label the actin cytoskeleton, with the cropped and zoomed ROI in 3(F).

**Figure 3.**
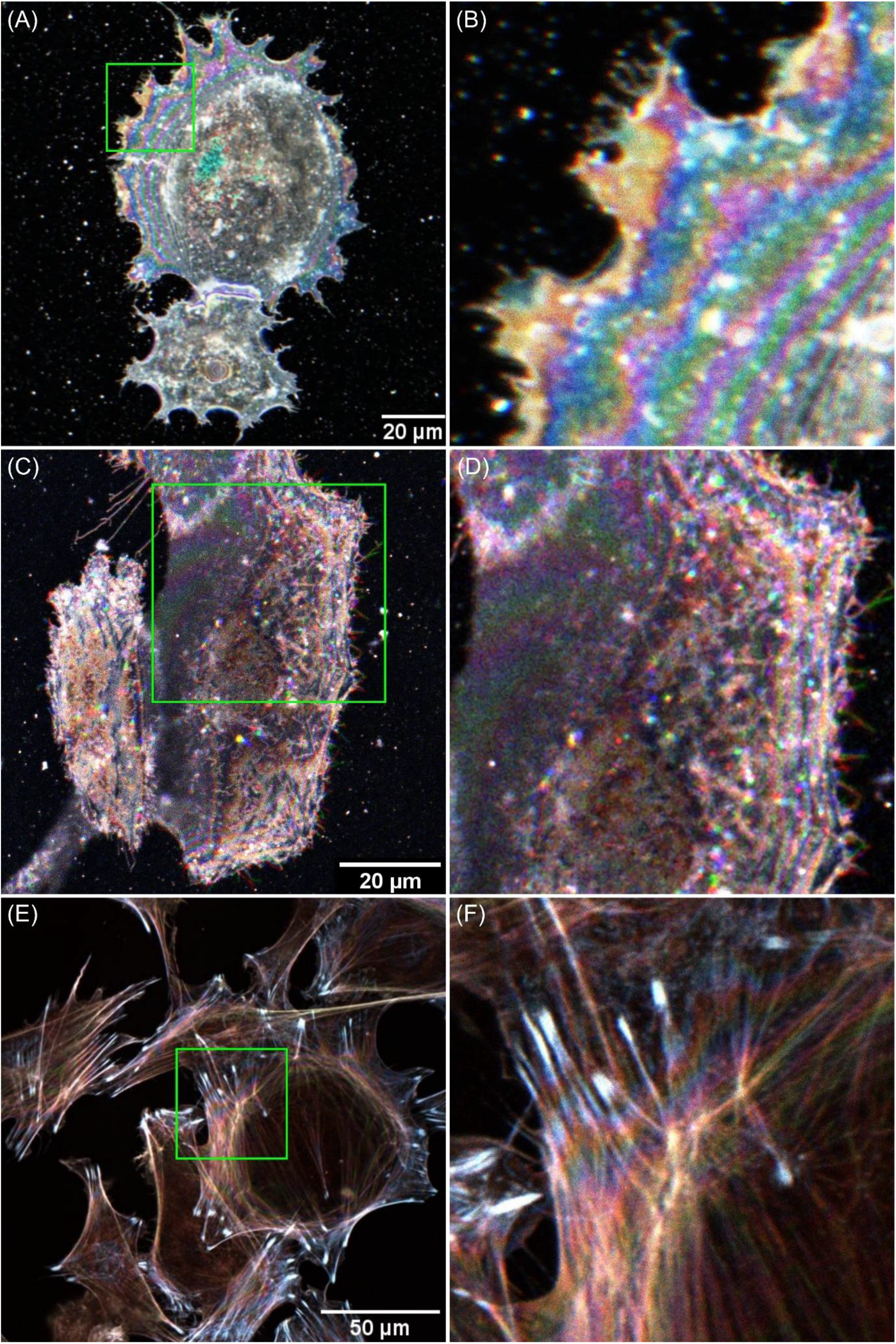
TartanSW cell images. (A) Live MCF-7 cells stained with the lipophilic stain Dil. Images were obtained with 2048 × 2048 pixels, and were averaged over 3 frames at a line speed of 100 Hz. The green box shows an ROI that is cropped, magnified and presented in (B). (C) shows live SKOV-3 cells labelled with the membrane dye Lipilight 560. Images were obtained with 2048 × 2048 pixels, and were averaged over 3 frames at a line speed of 100 Hz, with an emission detection bandwidth of 550-650 nm. The ROI indicated by the green box is shown in (D). (E) Fixed 3T3 cells prepared with rhodamine-conjugated phalloidin to label the actin cytoskeleton. Images were 2048 × 2048 pixels in size, and were averaged over 8 frames at a line speed of 100 Hz, with the indicated ROI shown in (F). For all images the 488 nm and 514 nm lines of an Argon laser and the 543 nm line of a Helium-neon laser were used for excitation of fluorescence, with an emission detection bandwidth of 550-650 nm.

In Figure 3(B) and 3(D), the pseudo-coloured anti-nodal planes are most clearly visible at the periphery of the cells, but they are difficult to discriminate in the centre of the cell. This is probably because of the highly-refractile cell organelles distorting the SW at the thicker parts of the cell. By manually counting the pseudo-coloured anti-nodal planes close to the edge of the cell, it is possible to measure the cell thickness up to around 3 μm from the basal membrane. However, it was not possible to reliably infer cell topology from the pseudo-coloured planes, even in the thin edge of the cell.

In Figure 3(F), where an actin label was used, the pseudo-coloured anti-nodal planes are still present, but they are less clearly visible. This is likely due to the complex morphology of the actin cytoskeleton. These pseudo-coloured anti-nodal planes do, however, persist deeper into the cell.

To show wider application of our TartanSW method we have also imaged other cell types and used alternatives to phalloidin stains. The data can be found in Supplementary Figure 1 and show similar results, demonstrating that the method works well with a range of spectrally different fluorophores and cell types. Supplementary Figure 1(A) shows fixed 1.1B4 cells labelled with rhodamine-conjugated phalloidin and excited with 488 nm, 514 nm and 543 nm laser lines. A cropped, magnified ROI of the image is shown in Supplementary Figure 1(B). Although an actin stain has been used, the filamentous network is so complex that it cannot be clearly resolved. However, due to the colour ordering in a TartanSW image, the shape of the apical cell membrane becomes clearly visible, and the presence of focal adhesions is revealed (white saturated regions, with examples of focal adhesions indicated by green arrows on Supplementary Figure 1B). Supplementary Figure 1(C) shows actin ruffles in fixed mouse PDAC (Pancreatic ductal adenocarcinoma) cells labelled with Alexa-488-conjugated phalloidin, with a siRNA knockdown of Aldolase A (Aldoa). Here, the sample was excited with 476 nm, 488nm and 496 nm laser lines. A cropped, magnified ROI is shown in Supplementary Figure 1(D). Similarly to 1(A), although an actin stain has been used, the filamentous network is so complex that it cannot be clearly resolved, and instead the overall cell topology is shown by the pseudo-coloured anti-nodal planes. However, Supplementary Figure 1(D) indicates that it is not simply the cell topology that can be seen. From the white saturated actin aggregate (which is an almost vertical thick band in Supplementary Figure 1(D), there are two different structures revealed by pseudo-coloured anti-nodal planes of opposite curvature. The brightest of these, which emits like a ‘C’ shape (indicated by a magenta arrow) with the curvature facing the linear saturated region, is likely from the apical membrane of the cell. There is a dimmer second ring pattern that emits like a ‘D’ shape (highlighted with a green arrow) with the curvature facing the linear saturated region. There are fewer anti-nodal planes present in this second feature, but, in combination, they indicate the presence of a tube-like structure that begins at the actin aggregate and proceeds into the cell body.

In order to confirm that these images are indeed created by SW phenomena, we performed controls in which the cells were adhering to coverslips instead of a mirror but otherwise applied the same imaging procedure. Example confocal images are shown in Supplementary Figure 2. As expected, we did not observe any pseudo-colour SW pattern when merging the three pseudo-coloured singleexcitation-wavelength images, due to the low reflectance at the mountant-to-coverslip interface.

We also applied the TartanSW method to live cells and were able to show that in principle the method can be used for long term live cell imaging, observing structural changes over time (Supplementary Video A & B). Supplementary Video A shows a movie of SKOV-3 cells labelled with the membrane dye Lipilight 560 and imaged over a period of 60 mins at 90 second intervals. As with the fixed cell data, the cell shows pseudo-coloured anti-nodal planes most clearly at the periphery of the cell. Over the duration of imaging, the intensity at the cell edge increases and becomes more saturated. This suggests accumulation of dye and possible restructuring of the membrane.

With cells expressing LifeAct GFP marking the actin (Supplementary Video B) we also observed anti-nodal planes, most prominently at the left-hand side of the cell close to the membrane which are moving over time.

## Discussion

The most important finding of our evaluation is the increase in sampling density in TartanSW microscopy compared to single-excitation-wavelength SW imaging. By sequentially exciting the same fluorophore with three different excitation wavelengths, the TartanSW method increased the sampling density from 50% to 98% over seven anti-nodal planes, with no notable change in axial or lateral resolution compared to single-excitation-wavelength SW microscopy.

The TartanSW method can be implemented into an existing confocal laser scanning microscope and does not require any adaptation to the hardware or image capture software. We have shown that the method is compatible with conventional fluorescent labels and photoproteins already used in fluorescence microscopy.

We had hoped that it may be possible to use the relative position of the pseudo-coloured anti-nodal planes to measure the shape of cell structure without the ambiguities that arise in single-excitation-wavelength SW microscopy. Specifically, we had hoped to be able to use the relative position of the pseudo-coloured anti-nodal planes to resolve two features in the axial direction with this method.

While it was possible to use *a priori* knowledge of the surface to reconstruct the TartanSW image of the lens specimen in 3D, unfortunately the fluorescently-tagged cell membrane and actin cytoskeleton proved too complex to be able to reliably resolve the differently coloured anti-nodal planes except in the thinnest part of the cell, close to the edge, so resolving adjacent structures in z with a resolution better than a confocal microscope did not prove feasible. However, it may be possible to overcome this limitation by using fluorescent dyes or photoproteins with a much broader excitation spectrum. A larger difference in wavelength between the different excitation wavelengths should, in theory, allow simpler indexing of the anti-nodal planes by colour order. We were unable to measure the effect of the excitation spectrum bandwidth on indexing by colour because the dyes and photoproteins that we used had a similar bandwidth, but alternative dyes may offer some improvement in the future.

We used an upright confocal microscope in our evaluation, but we also explored the application of TartanSW on an inverted microscope. For that, we prepared the cells on mirrors as described previously, and placed them into coverslip-bottom dishes, with the mirror surface facing the coverslip. We tried dry, water immersion and oil immersion objective lenses, but with an inverted microscope no SWs could be detected. We believe this is due to the mass of the mirror compressing the cell volume significantly onto the coverslip such that the specimen became almost two dimensional. As such, we would recommend TartanSW only to be used with an upright microscope.

Further advantages of TartanSW for three-dimensional imaging are the increased axial resolution and high speed compared to CLSM. The axial resolution of the CLSM image is determined by the PSF of the microscope, whereas the axial resolution of the TartanSW image depends on the thickness of the anti-nodal planes. For the 40x/0.8 N.A. HCX APO L U-V-I lens used in our cell imaging work and at a wavelength of 543 nm the theoretical axial resolution in standard light microscopy was calculated to be 1.7 μm, whereas, as shown in Table 1, the anti-nodal plane thickness in the TartanSW image was 139 nm, representing a resolution improvement of more than an order of magnitude. The speed advantage was also considerable. For a 4096 × 4096 pixel image at a line speed of 100 Hz and a frame average of 8 it takes approximately 10 mins to acquire one TartanSW image with three excitation wavelengths. For the same imaging parameters and using a z step size of 420 nm for Nyquist sampling in the axial dimension with the N.A. = 0.8 objective lens, a CLSM z-stack of the same volume takes over 2 hours because of the need to move the specimen or objective lens in the z-direction for three-dimensional imaging. TartanSW may therefore offer reduced photobleaching and phototoxicity compared to CLSM, which has advantages for long-term three-dimensional cell imaging. However, the need for sequential laser excitation in TartanSW may prove limiting for live cell studies where cell structures move or change rapidly. This is evident in Figure 3(D), where some of the dye spots are not singular and white in the RGB merge but instead are multi-colour and consist of several spots very close together. Since these are not present in the fixed cell data and they do not occur with uniform directionality in the image, we do not attribute this to chromatic aberration, or to specimen or microscope stage drift. Instead, we expect that these changes in spot location arise from dynamic changes within the cell.

Although our TartanSW results show promise for cell imaging studies, as with single-excitation-wavelength SW microscopy, it is currently very difficult to make a three-dimensional reconstruction of the cell. We have previously used the theoretical values of anti-nodal spacing with single-excitation-wavelength SW microscopy data to create a three-dimensional reconstruction of the basal membrane of a red blood cell, but even for one SW this is computationally demanding and there is uncertainty in the approximations involved [24]. In TartanSW, the third-dimensional information is present as contour lines arising from the anti-nodal planes within a two-dimensional image, and the relative heights of the contour lines are unknown. As described above, it is possible that the use of fluorescent dyes with a broader excitation spectrum above will separate the anti-nodal planes further, aiding indexing by colour. If successful, this would also likely help to achieve the sought-after three-dimensional reconstruction of a SW cell image dataset.

## Methods

### Non-biological test specimen preparation

An uncoated silica plano-convex lens with a focal length of 48mm (Edmund Optics Ltd, York, UK) was prepared as described previously [24].

### Biological specimen preparation

The cell lines utilised in this study were breast cancer cells (MCF-7, (ATCC^®^ HTB-22)), ovarian cancer cells (SKOV-3, ATCC^®^ HTB-77), and pancreatic Beta cells: PANC-1 hybrid (1.1B4), fibroblast (3T3, (ATCC^®^ CRL-1658)) and pancreatic ductal adenocarcinoma cells (PDAC, ATCC^®^ CRL-1469). PDAC cells from KPC mice [26] were a gift from Saadia Karim, Cancer Research UK Beatson Institute [27].

Cells were grown in vented capped tissue culture flasks containing DMEM (Gibco 10567-014, Thermo Fisher, Paisley, UK) supplemented with 10% FBS (Labtech, Heathfield, UK), 1% Penicillin Streptomycin (Gibco, Thermo Fisher, Paisley, UK) and 0.6% L-glutamine (Thermo Fisher, Paisley, UK), and were incubated at a temperature of 37 °C in a 5% CO_2_ humidified cell incubator (Heracell VIOS CO_2_, Thermo Fisher, Paisley, UK).

For TartanSW imaging, individual cell lines were seeded onto separate mirrors (TFA-20C03-10, Laser 2000, Huntingdon, UK) in 6 well plates at the desired density and incubated for 24 h at 37 °C / 5% CO_2_ to allow adherence to the mirror. We chose these mirrors for their high flatness (λ/10) which we thought may be needed when using an objective lens with a small depth of field. However, when imaging with an objective with a larger depth of field, flatness of mirror is less of a concern. Prior to imaging, the cells were either fixed (as described below) or stained for live cell imaging. For control cell specimens, the cells were seeded onto Type 1.5, 18 mm diameter circular borosilicate glass coverslips (VWR, Lutterworth, UK) and they were otherwise treated identically to the cell specimens grown on mirrors. Cells adhering to mirrors or coverslips were washed once to remove culture medium with the respective buffer before staining, at least three mirrors or coverslips with cells were prepared.

For DiI staining, washed cells were transferred into fresh 4% bovine serum albumin (BSA, Sigma Aldrich, Dorset, UK) in 1x phosphate buffered saline (PBS, Thermo Fisher, Paisley, UK), 20 μl of 1 mg/ml DiI stock (Thermo Fisher, Paisley, UK) was prepared in DMSO (Sigma Aldrich, Dorset, UK) and added to 4 ml of buffer, followed by an incubation for 1 hour at 37°C. The cells were washed twice in 1x PBS for 5 min, then placed in 4% BSA in PBS for imaging.

For Lipilight staining (Idylle, Paris, France), after twice washing the cells with PBS (1x PBS, 5 min) 50 nM of Lipilight 560 was added to the cells. After 3 minutes incubation, the sample was imaged immediately.

For phalloidin-based stains, (Rhodamine-conjugated phalloidin, and Alexa488-conjugated phalloidin, all from Thermo Fisher, Paisley, UK), cells were washed twice with PBS (1x PBS, 5 min) and then fixed in 4% PFA (Paraformaldehyde, Sigma Aldrich, Dorset, UK) for 10 min, washed twice in PBS (1x PBS, 5 min), permeabilised in 0.1% Triton X/PBS for 5 min, washed twice in PBS (1x PBS, 5 min) and were then blocked with 1% BSA in PBS for 30 min. This was followed by an incubation in a 1/2000 phalloidin-based stain in 1% BSA/PBS buffer for 20 min in light-tight conditions, after which the specimen was washed twice (1x PBS, 30 s) and stored in PBS at 4°C until imaged.

The 3T3 cell line was transfected with a LifeAct GFP construct (pEGFP-N1-LifeAct-*FcoR*I-*BamH*I) using a Polyplus jetOPTIMUS transfection reagent (Polyplus transfection, Illkirch, France) following the manufacturer’s instructions.

Knockdown of Aldolase A (Aldoa) in mouse PDAC cells was performed using Lipofectamine RNAiMax (ThermoFisher Scientific, 13778150) and Mm Aldoa 2 FlexiTube siRNA (Qiagen, SI00896238) according to the manufacturer’s instructions.

### TartanSW imaging

All the SW imaging described in this paper was done in laser scanning confocal mode, using a Leica SP5 DM600 microscope (Leica Microsystems, Wetzlar, Germany). We used a 5x/0.15 N.A. HCX PL FLUOTAR DRY objective for imaging the non-biological test specimen, which was air-mounted, and a water dipping objective, 40x/0.8 N.A. HCX APO L U-V-I, for cell imaging. Cell specimens were mounted in PBS (n=1.34), and all images were acquired with sequential laser excitation. The confocal aperture was set to 1 Airy unit for all images. Opening the pinhole did not change the result except to increase the fluorescence intensity signal of the image

For imaging of the non-biological lens test specimen, live SKOV-3 cells labelled with Lipilight 560 and live MCF-7 cells labelled with DiI, the 488 nm and 514 nm lines of an Argon laser and the 543 nm line of a Helium-neon laser were used for excitation of fluorescence. The SKOV-3 cells were imaged over 1 hour with 90 s time lapse intervals. Images were 2048 × 2048 pixels, and were averaged over 3 frames at a line speed of 100 Hz, with an emission detection bandwidth of 550-650 nm. For imaging of fixed 3T3 cells and fixed 1.1B4 cells labelled with Rhodamine-conjugated phalloidin, the 488 nm and 514 nm lines of an Argon laser and the 543 nm line of a Helium-neon laser were used for excitation of fluorescence. Images were 2048 × 2048 pixels, and were averaged over 8 frames at a line speed of 100 Hz, with an emission detection bandwidth of 550-650 nm. Fixed PDAC cells labelled with Alexa-488-conjugated phalloidin were excited with the 476 nm, 488nm and 496 nm lines from an Argon laser. Images were 4096 × 4096 pixels, and were averaged over 8 frames at a line speed of 100 Hz, with an emission detection bandwidth of 505-600 nm. For live GFP LifeAct transfected 3T3 cells, the excitation came from the 476 nm, 488 nm and 496 nm lines of an Argon laser. The image size was 1024 × 1024 pixels, and images were averaged over 3 frames at a line speed of 100 Hz. The emission was detected between 520-625 nm. Images were obtained at 48 second intervals.

In order to confirm that the presence of anti-nodal planes and colour ordering were indeed a result of the TartanSW method, we also imaged cells adhered to coverslips instead of grown on a mirror and imaged them with standard confocal microscopy, using the same imaging parameters as described above.

### Image processing

The sequentially acquired images of the specimen for the different excitation wavelengths were contrast adjusted using the ‘Auto’ Brightness/Contrast function in FIJI [28] and merged using the pseudo colours red, green and blue, with red being used for the longest excitation wavelength and blue for the shortest excitation wavelength of the individual imaging experiment.

In FIJI, the non-biological specimen image data was cropped and a Gaussian blur *of σ* = 2 was applied to remove noise. In MATLAB, a radial line profile was taken from the centre of the TartanSW image outwards and, by using Eqn. 1 (Supplementary File 1, SF1)

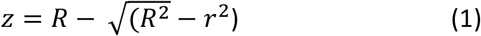

where *z* is the height from the mirror, *R* is the radius of the curvature, and *r* is the radial distance, the radial distance was translated to axial height. As a result, an intensity profile was plotted against axial height (relative to the mirror surface) over the range of *z* = 5 μm. An axial range of 5 μm was selected to characterise the spectral signature over a large axial distance. The information gap was calculated by using the *findpeaks* function in MATLAB, where only peaks with a normalised intensity of above 0.1 and a peak separation of 100 nm were included in the analysis. From the detected peaks, the measured full-width at half-maximum (FWHM) was extracted and the minimum FWHM boundary from the shortest wavelength (488 nm) and the maximum FWHM of the longest wavelength (543 nm) were extracted. This was repeated for seven anti-nodal planes, as to keep the comparison consistent across the SW images.

To determine whether our proposed TartanSW method would increase the sampling density, the seven 488 nm and 543 nm FWHM boundary values were subtracted and the result for each boundary was added together to determine the total contribution of the TartanSW multi-excitation method. Next, the total contribution was subtracted from the total distance over the eighth nodal plane. The percentage of information gain between the TartanSW image and a single wavelength SW microscopy image at 543 nm was used to determine the increase in sampling density resulting from the introduction of two additional SWs for excitation. The fill factor was calculated by a summation of the contribution from all wavelengths, where only the values greater than the FWHM of the standing wave antinode were counted. Once these values were summed, the fill factor was calculated as a percentage of the total contribution of the filled regions, with respect to the total height which corresponded to the distance from 0 to the 8^th^ nodal plane of the 543nm standing wave. The custom-written MATLAB code to perform this measurement is provided as Supplementary File 1 (SF1 and SF2).

MATLAB was used to make a 3D reconstruction of the lens specimen imaged using the TartanSW method. Firstly, the image was inputted into MATLAB using the imread function. From the TartanSW image, two matrices were generated, using MATLABs meshgrid function, which corresponded to the × and y positions of each pixel. Next, the × and y matrices were then multiplied by the image scale factor of 1.39 to convert the pixel values to micrometer. The radial distance, r, was then calculated from the × and y pixel distance matrices using Pythagoras theorem. The axial height of each pixel in micrometer was calculated from Eqn. 1 using the radial distance r, and the radius of the curvature, R, of the lens specimen. Lastly, a 3D reconstruction of the lens specimen was created using the x, y and z pixel values, and an RGB colour map obtained from the TartanSW image, using MATLABs scatter3 function. The custom-written MATLAB code to perform this measurement is provided as Supplementary File 1 (SF3).

## Acknowledgements

This work was supported by the Biotechnology and Biological Sciences Research Council, grant number BB/P02565X/1. Lisa Kölln and Katrina Wesencraft were supported by the Medical Research Council and Engineering and Physical Sciences Research Council Centre for Doctoral Training in Optical Medical Imaging, grant number EP/L016559/1. Peter W. Tinning, Gillian Robb and Gail McConnell were partly supported by the Medical Research Council, grant number MR/K015583/1. Nikki Paul was supported by Medical Research Council grant MR/R017247/1 and Cancer Research UK grants A17196 and A24452.

We thank Jan Schniete for his support and troubleshooting for the MATLAB code and helpful discussions.We thank Maddy Parsons (King’s College London, UK) for the gift of the LifeAct GFP construct pEGFP-N1-Lifeact-*EcoR*I-*BamH*I.

## Author contributions

JKS, PWT, TB, GM contributed to the writing of the manuscript. GM & TB obtained the funding and conceived the project. RS wrote the MATLAB code and simulation of TartanSW data. JKS, PWT, RS, GR, KW, NP prepared sample. JKS, RS, GR, LK, KW, NP, GM carried out the imaging of the samples. JKS, PWT, RS, LK, GM carried out the image analysis

## Data Availability

All data generated or analysed during this study are included in this published article (and its Supplementary Information files) and are also deposited here 10.15129/fd3622cc-bede-402d-8170-50367f736e41

## Supplementary Files and Figures

**Supplementary Figure 1.**
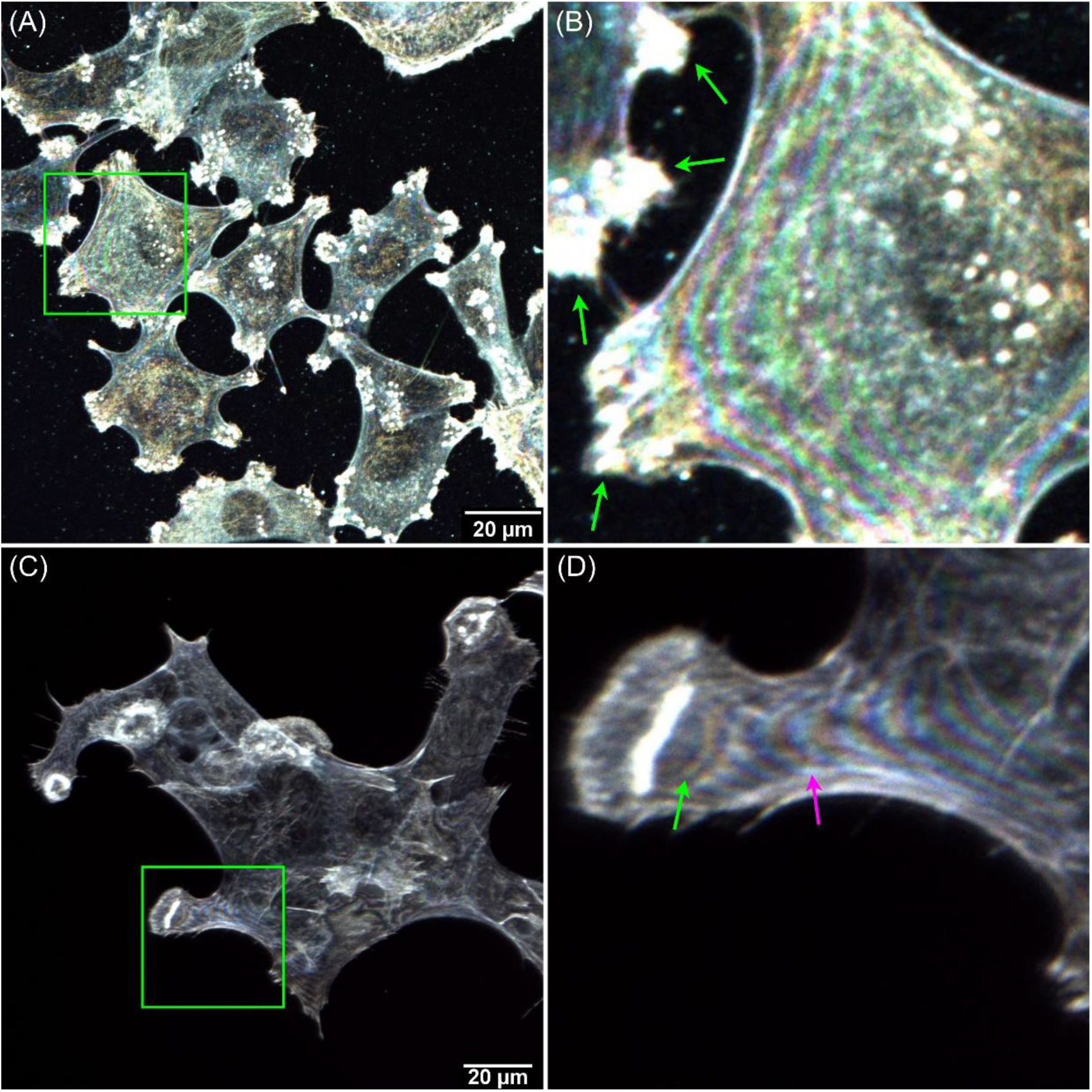
Examples of TartanSW imaging of the actin network in fixed cell lines. (A) Pancreatic beta cell line 1.1B4 labelled with rhodamine-conjugated phalloidin. The 488 nm and 514 nm lines of an Argon laser and the 543 nm line of a Helium-Neon laser were used for excitation of fluorescence. Images were taken with 2048 × 2048 pixels, and were averaged over 8 frames at a line speed of 100 Hz, with an emission detection bandwidth of 550-650 nm. A green box indicates an ROI, which is cropped and magnified and shown in (B), where focal adhesions are indicated with green arrows. (C) Actin ruffles in fixed mouse PDAC (Pancreatic ductal adenocarcinoma) cells labelled with Alexa-488-conjugated phalloidin with a siRNA knockdown of Aldolase A (Aldoa). Fluorescence was excited with the 476 nm, 488nm and 496 nm lines from an Argon laser. Images were 4096 × 4096 pixels, and were averaged over 8 frames at a line speed of 100 Hz, with an emission detection bandwidth of 505-600 nm. A green box indicates an ROI, which is cropped and magnified and shown in (D). The magenta and green arrows highlight the ‘C’ and ‘D’ shaped curvature of two standing waves that indicate the presence of a tube-like structure beginning at the actin aggregate and proceeding into the cell body.

**Supplementary Figure 2.**
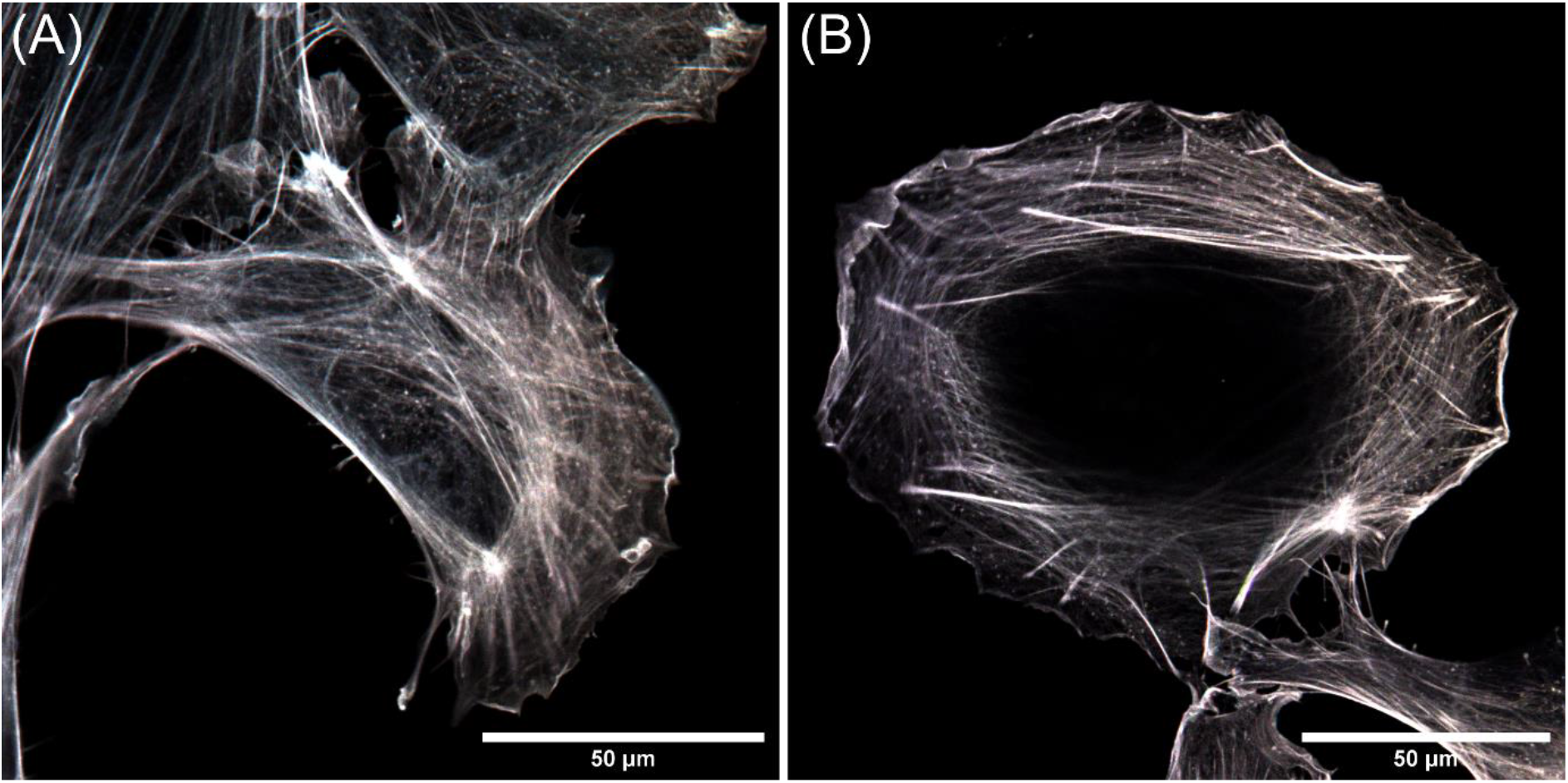
Control studies confirm that the mirror is an essential part of the TartanSW method. Fixed 3T3 cells were grown on coverslips instead of a mirror, and were stained with rhodamine-conjugated phalloidin as control specimens. These preparations were imaged with the same parameters as described in Figure 3(E) and (F). (A) and (B) show two example RGB colour-merge images of cells. No pseudo-coloured anti-nodal planes are visible in the data.

**Supplementary Figure 3.**
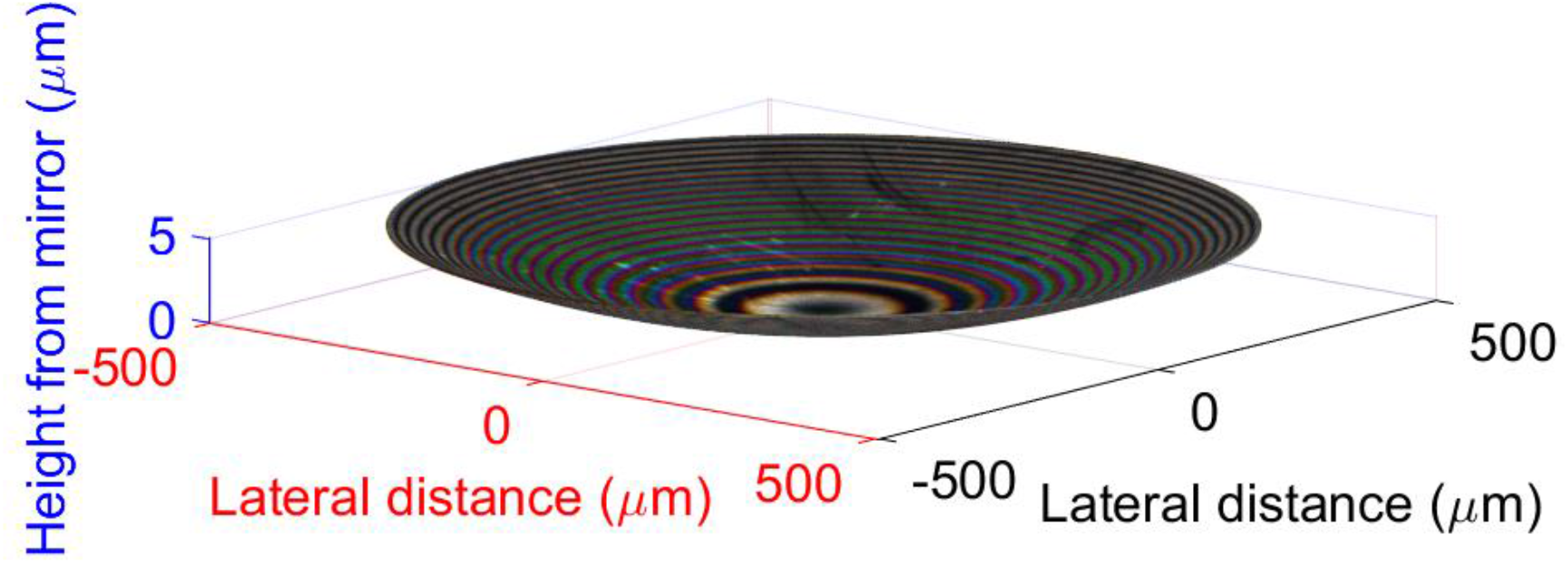
3D reconstruction a lens specimen imaged using the TartanSW method from a 2D image using a custom written MATLAB code (see SF3)by generating two matrices containing the × and y positions of each pixel which where than converted to the corresponding height by a conversion factor. The radial distance, r, was then calculated from the × and y pixel distance matrices using Pythagoras theorem. The axial height of each pixel in micrometer was calculated from Eqn. 1 using the radial distance r, and the radius of the curvature, R, of the lens specimen. Lastly, a 3D reconstruction of the lens specimen was created using the x, y and z pixel values, and an RGB colour map obtained from the TartanSW image, using MATLABs scatter3 function

### Supplementary videos - Live cell imaging using TartanSW

The videos available here: 10.15129/fd3622cc-bede-402d-8170-50367f736e41 named as “Supplementary Video A SKOV-3 Lipilight_560.avi” and “Supplementary Video B 3T3 GFP_Lifeact.avi”

(A) Live SKOV-3 cells labelled with the membrane dye Lipilight 560. Images were obtained with 2048 × 2048 pixels over a period of 1 hour at 90 second intervals. A frame average of three was applied at a line speed of 100 Hz. The 488 nm and 514 nm lines of an Argon laser and the 543 nm line of a HeliumNeon laser were used for excitation of fluorescence, which was detected over an emission bandwidth of 550-650 nm.

(B) GFP LifeAct transfected 3T3 cells. Images were obtained with 1024 × 1024 pixels over a period of 40 minutes at 48 seconds intervals. A frame average of three was applied at a line speed of 100 Hz. The 476 nm, 488nm and 496 nm lines from an Argon laser were used for excitation and fluorescence was detected over an emission bandwidth of 520-625 nm.

### Supplementary Files: Matlab Code

SF1 Matlab Code I: TartanLensanalysis.m

SF2 Matlab Code II–radialavg.m

SF3 Matlab Code III–3Dreconstruction_lens.m

